# Fluorogenic probe for fast 3D whole-cell DNA-PAINT

**DOI:** 10.1101/2020.04.29.066886

**Authors:** Kenny KH Chung, Zhao Zhang, Phylicia Kidd, Yongdeng Zhang, Nathan D Williams, Bennett Rollins, Yang Yang, Chenxiang Lin, David Baddeley, Joerg Bewersdorf

## Abstract

DNA-PAINT is an increasingly popular super-resolution microscopy method that can acquire high-fidelity images at nanometer resolution. It suffers, however, from high background and very slow imaging speed, both of which can be attributed to the presence of unbound fluorophores in solution. We present a fluorogenic DNA-PAINT probe that solves these problems and demonstrate 3D imaging without the need for optical sectioning and a 26-fold increase in imaging speed over regular DNA-PAINT.

## Main Text

Single-molecule localization microscopy (SMLM) has become an invaluable tool in biological research, revealing subcellular structure and function at ten or more times the resolution of conventional fluorescence microscopy ^1^. Among these techniques, DNA-based points accumulation for imaging nanoscale topography (DNA-PAINT) microscopy is becoming increasingly popular because of the high-quality images that can be obtained with relative ease ^2–5^. DNA-PAINT, in contrast to other SMLM techniques, does not rely on light to switch fluorescent molecules between ‘visible’ and ‘invisible’ states ^2^. Instead, it takes advantage of transient DNA binding to achieve the same switching or ‘blinking’ effect. Single-stranded DNA (ssDNA) ‘docking strands’ attached to the biological structure of interest are individually highlighted as they capture an ‘imager probe’, a complementary ssDNA molecule linked to a fluorophore, from the diffusing pool.

There are several advantages to this approach: (1) Binding-based blinking frequency is proportional to the imager probe concentration and is therefore easily adjustable (see **Suppl. Note 1**) ^2^. (2) Fluorophores and buffers are not limited to those suitable for photophysical blinking but can be chosen to allow for maximum blink brightness and thereby best localization precision ^2, 6^. (3) DNA-PAINT is robust against photobleaching as bleached probes are replenished from the large reservoir of probes in solution ^5^. (4) DNA-PAINT can take advantage of the versatile toolbox for *in-vitro* DNA technology, for example creating libraries of fluorescence in situ hybridization (FISH) probes for OligoDNA-PAINT ^4^, use of photoreactive nucleosides in Action-PAINT ^7^, or click chemistry reactions for labeling ^8^.

However, the use of transient chemical binding-based switching is also responsible for the greatest weakness of DNA-PAINT: unbound imager probes contribute massive amounts of background fluorescence which can drown out the signal peaks of individual bound probe molecules. To minimize background, early DNA-PAINT realizations relied on low probe concentrations and the use of total internal reflection fluorescence (TIRF) to minimize illumination volume ^9^. Consequently, DNA-PAINT was limited to imaging regions within a few 100 nm of the coverslip. Using a spinning disk confocal microscope, regions deeper in a cell could be imaged, but at the cost of a compromise in resolution, caused by lower detected photon numbers per blinking event ^10^. The constraint on probe concentration to typically less than 5 nM also limits the number of blinking events per second, leading to slow imaging speeds. This problem can be mitigated to some extent by increasing the rate constant of an imager probe binding a docking strand, *k*_*ON*_, as recently demonstrated ^11, 12^. While these publications could show super-resolution images acquired in minutes instead of hours, they still required TIRF to keep background low.

None of these approaches, however, solves another background-related problem: at high camera frame rates, as required for fast imaging, exposures are so short that unbound molecules are no longer blurred by diffusion. These molecules will appear as local fluctuations in background and, at worst, erroneous binding events. To address both background-related issues requires a reduction in the molecular brightness of the diffusing unbound probe molecules. Recently, two groups introduced Förster resonance energy transfer (FRET)-based DNA-PAINT to achieve this effect ^13–15^. Donor and acceptor fluorophores were conjugated to separate imager probes; only when they coincidentally bind to the same docking strand will donor excitation lead to acceptor emission. Unfortunately, the demonstrated localization precision was substantially worse than for regular DNA-PAINT, failing the ‘gold standard’ of resolving the hollow center of labeled microtubules. This reduction in image quality can be attributed to FRET DNA-PAINT being fundamentally limited by the trade-off between maximizing energy transfer efficiency for bright blinking events and minimizing excitation and emission crosstalk to reduce background.

High background fluorescence therefore remains a substantial problem in DNA-PAINT and severely limits its application: it has not yet been possible to image whole cells in 3D using DNA-PAINT without optical sectioning. Even with optical sectioning, compromises are made between quality and speed, requiring either imaging for hours at low probe concentration to acquire high-quality DNA-PAINT images or for minutes at high probe concentration at the risk of undersampling, resolution loss, and the introduction of artifacts from too many molecules blinking at once and high, inhomogeneous background.

Here we introduce a new type of DNA-PAINT probe that is highly fluorogenic, i.e. is dark when in solution and bright when bound to a docking strand (Fig. 1A iv). This feature enables us to perform high-quality DNA-PAINT imaging without optical sectioning. It also allows us to substantially increase the probe concentration and record blinking events at 100 frames per second (fps), with negligible compromises in localization precision.

**Figure 1.**
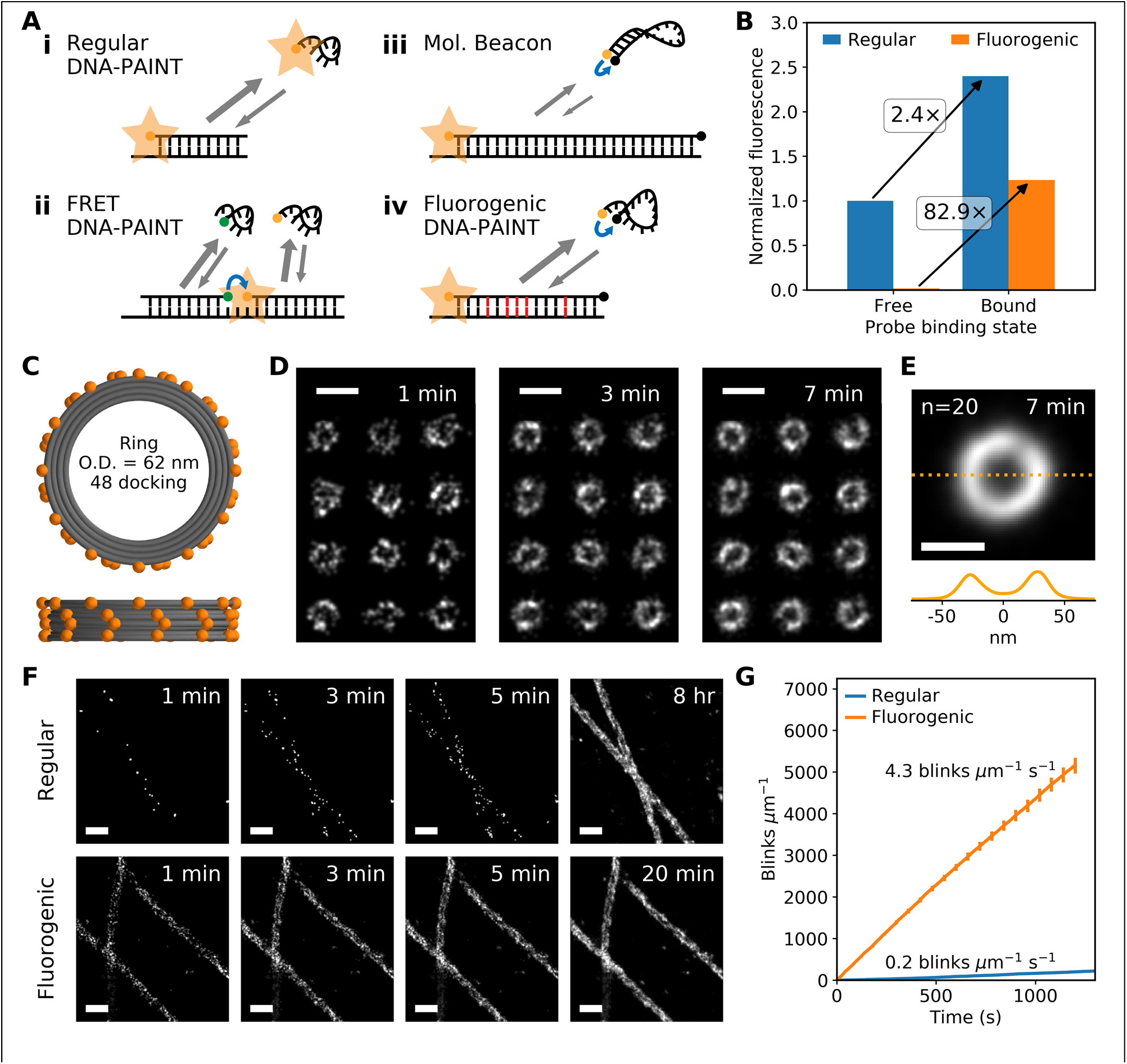
Comparison of regular and fluorogenic DNA-PAINT probes. **(A)** Comparison of different DNA-PAINT imager probe / docking strand systems: **(i)** Imager probe for regular DNA-PAINT is always fluorescent and contributes to high background. **(ii)** In FRET DNA-PAINT, acceptor fluorescence is only observed when both donor- and acceptor-conjugated probes are bound to the docking strand. (Energy transfer indicated by blue arrows.) **(iii)** Regular molecular beacons are fluorogenic but their binding on- and off-rates are too slow for DNA-PAINT. **(iv)** Our DNA-PAINT probe is fluorogenic and has fast binding kinetics due to the absence of a stem secondary structure and mismatches between the probe and docking sequences (shown in red). **(B)** Comparison of fluorescence between a regular imager probe (Cy3B) and our fluorogenic imager probe (Cy3B and BHQ-2) in solution. In their unbound ssDNA state, the fluorogenic probe is less than 1.5% as bright as the regular probe. On binding to their complementary docking strand, its fluorescence increases 83-fold, compared to a 2.4-fold increase for the regular probe. **(C)** Top and front schematic view of the DNA origami nanostructure, with 48 docking strands attached to the 62-nm ring. **(D)** Fast fluorogenic DNA-PAINT TIRF imaging ([probe] = 250 nM; frame rate = 100 Hz) of origami rings (12 rings shown). Scale bar is 100 nm. **(E)** Average of 20 rings measuring a diameter of 60.5 ± 0.5 nm and displaying ~9% multi-emitter artifacts (**Suppl. Fig. 3**). Scale bar is 50 nm. **(F)** Qualitative comparison of regular versus fluorogenic DNAPAINT TIRF-imaging of fixed microtubules (regular: [probe] = 100 pM, frame rate = 4 Hz; fluorogenic: [probe] = 20 nM, framerate = 100 Hz). Scale bar is 200 nm. **(G)** Quantitative comparison showing that blinking data is acquired 25.8 times faster with the fluorogenic probe (quantified by the number of blinks per length of microtubule per unit of time; regular 0.17 ± 0.01 μm^−1^ s^−1^, n_ROI_ = 8, fast 4.30 ± 0.16 μm^−1^ s^−1^, n_ROI_ = 10). Averaged statistics presented as mean ± SEM.

Our new approach is based on the conceptual realization that using FRET for the *suppression of background rather than the induction of signal* avoids the weakness of FRET DNA-PAINT. Using a non-fluorescent quencher as an acceptor and measuring fluorescence from the donor eliminates cross-talk, allowing us to maximize quenching efficiency. We can use dedicated quenchers, which are very efficient FRET acceptors due to their high extinction coefficient and broad absorption spectrum, and can also utilize non-FRET based quenching mechanisms ^16^. Since donor fluorescence is detected in the absence of energy transfer to the acceptor, the donor can still be chosen based on its photon count to achieve the best localization precision. With this design, the donor and acceptor do not act as coincidence detectors and can be conjugated to the two terminals of the same imager probe, which preserves the photobleaching resistance of DNA-PAINT and simplifies probe kinetics design (Fig. 1A iv). Instead, our design relies on the changes in conformation of the imager probe upon binding to the docking strand. Unbound ssDNA is highly flexible which causes the fluorophore and quencher to be in close proximity and results in strong quenching. In contrast, when complemented by the docking strand, the resulting dsDNA takes the form of a rigid double helix and the terminals are spatially separated to minimize quenching. The success of this concept has been shown with molecular beacons, hairpin ssDNA structures which are quenched in solution but unfold when bound to its target nucleic acid sequences ^17^. However, the slow kinetics of these probes (*k*_*ON*_ < 10^4^ M^−1^s^−1^, off-rate ≪ 1 s^−1^) make them unsuitable for DNA-PAINT (*k*_*ON*_ > 10^5^ M^−1^s^−1^, off-rate > ~1 s^−1^) ^18^. We set out to rationally design a fluorogenic imager probe/docking strand system for DNA-PAINT by prioritizing four key properties: (1) To minimize background, freely diffusing probes should be as dark as possible, i.e. they should have a low molecular brightness; (2) *k*_*ON*_ should be comparable to (or higher than) regular DNA-PAINT to avoid excessively high probe concentrations and background; (3) unbinding should occur quickly (off-rate should be high) to allow for maximum imaging speed (see **Suppl. Note 2** and **Suppl. Fig. 2**); (4) bound probes need to unquench effectively to not compromise localization precision of the blinks.

**Figure 2.**
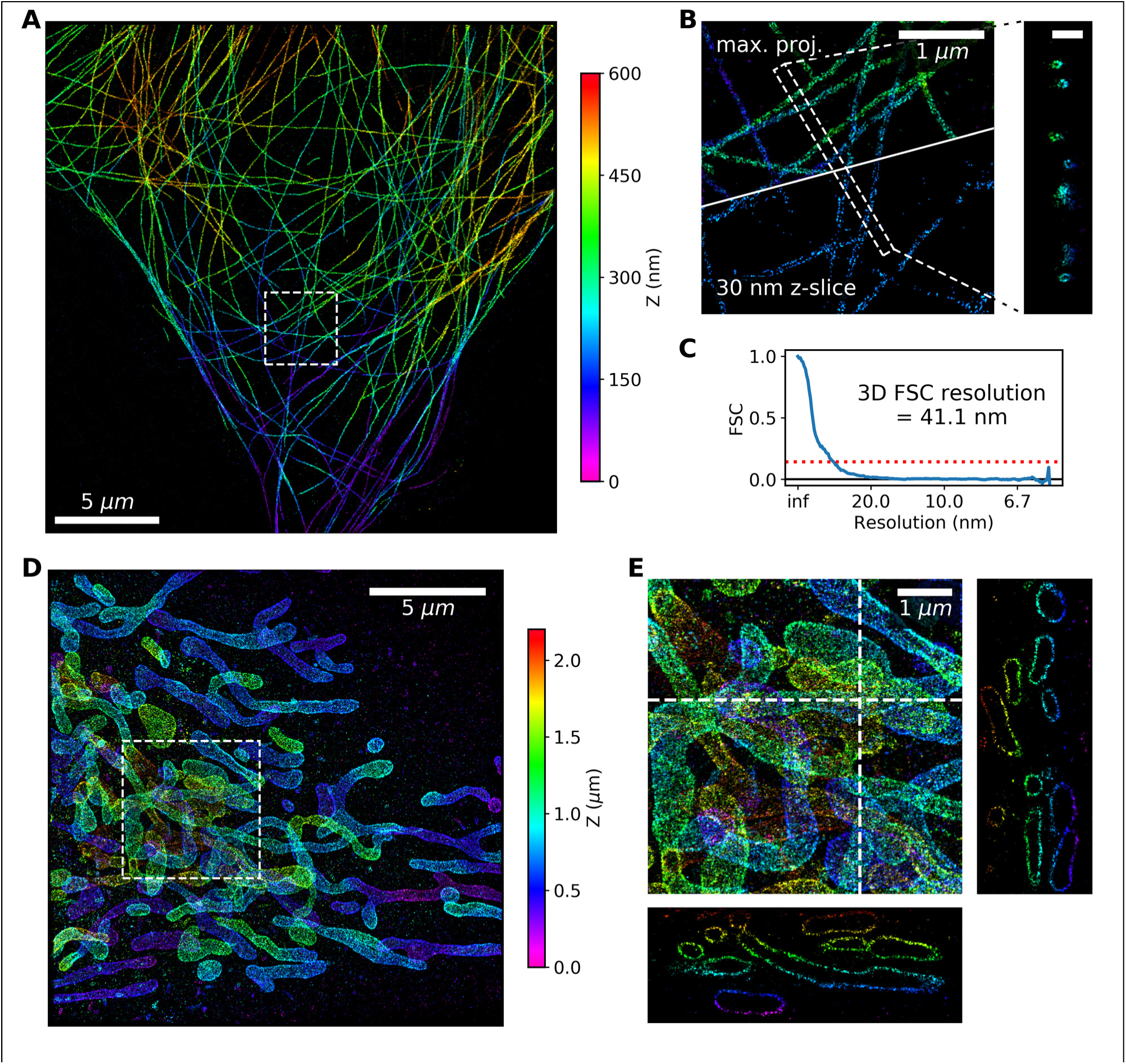
Fluorogenic probes enable fast 3D DNA-PAINT imaging without optical sectioning. **(A-C)** Fast 3D fluorogenic DNA-PAINT imaging ([probe] = 10 nM; 100 Hz; 10 min) of immunolabeled microtubules in COS-7 cells without optical sectioning under widefield illumination. **(A)** Overview image. **(B)** Zoomed-in view of the dashed box in (A). The top represents a maximum projection image, the bottom a 30-nm thick slice. (A). Hollow microtubules can be seen in the slice view as well as in the Z cross-section displayed on the right (scale bar: 250 nm) **(C)** Fourier shell correlation (FSC) with a threshold of 0.143 measures the 3D resolution at 41.1 nm. **(D, E)** Fast 3D whole-cell fluorogenic DNA-PAINT imaging of mitochondria in U-2 OS cells overexpressing GFP-OMP25 recorded with a 4Pi-SMS microscope ([probe] = 2 nM; 100 Hz; 1 hr). Regular DNA-PAINT is incompatible with 4Pi-SMS due to its lack of optical sectioning. **(D)** Zprojection of data collected from 4 optical planes covering a depth range of 2 μm. **(E)** Magnified region from **(D)** with XZ and YZ cross-sections.

The success of our system required solving the conflict between the need for large distances between the fluorophore and quencher to avoid quenching of the bound probe and the desire for high off-rates to allow for fast imaging. The former demands a long stretch of rigid probe of no less than about 15 bases (0.34 nm per base along helical axis versus a Förster radius between the quencher and fluorophore of ~6 nm) ^19^. In contrast, the latter needs 10 or less complementary base pairs between the imager probe and docking strand. We solved what superficially appears to be a conflict by designing docking strand sequences with internal mismatches against a long 15-bases imager probe to destabilize the binding for a faster off-rate.

Additionally, we needed to avoid stabilizing the free imager probes in their quenched form, which can happen for example by a self-complementary sequence or ‘stem’ in the probe, since this effect would render *k*_*ON*_ too slow ^11^. We hypothesized that even in the absence of a stem, ssDNA without secondary structure would be sufficiently dynamic and flexible to provide strong quenching in the unbound state. Based on these considerations, we designed a 15-bases long probe and screened a number of docking strands before choosing the probe/docking strand pair used in the study.

We first characterized the fluorogenicity of our imager probe in solution. In its freely diffusing unbound state, our probe is less than 1.5% as bright as a regular DNA-PAINT probe (Fig. 1B). On binding with its fully complementary sequence, we observed a 83-fold increase in fluorescence. Interestingly, we also measured a weak fluorogenic effect of a 2.4-fold increase for the regular probe. Overall, our new imager probe was still 35 times more fluorogenic than the regular DNA-PAINT probe.

We first imaged DNA origami nanostructures because the position and number of the docking strands can be precisely defined. The chosen ring shaped origami was originally intended as a mimic for nuclear pore complexes^20^. and has an outer diameter of 62 nm to which 48 docking strands were added. This relatively low density of docking strands compared to typical antibody labels in cells makes DNA-PAINT imaging very inefficient unless the imager probe concentration is high enough to lead to substantial binding frequencies. It therefore represents an excellent test of the performance of DNA-PAINT under conditions where the background is high. The ring shape provides an easy and visual way to assess the achievable resolution as well as blinking artifacts. Imaging with an imager probe concentration of 250 nM, around 50 times higher than regular DNA-PAINT, and a camera frame rate of 100 Hz with TIRF illumination, we could resolve the rings within 1 minute (Fig. 1D). The ring diameter was measured to be marginally less than the expected size (60.5 ± 0.5 nm (mean ± SEM), n=20). Fourier Ring Correlation (FRC) analysis yielded an excellent resolution of 17.8 nm (see **Methods**). The hollow centers of the rings could be easily resolved in individual rings as well as in an average of 20 rings (Fig. 1E) confirming that the frequency of multi-emitter artifacts is low (~9%; **Suppl. Fig. 3**).

For quantification of imaging speed and resolution with a biological sample that features a large number of docking strands per diffraction-limited area, we compared 2D DNA-PAINT TIRF images of immunolabeled microtubules in COS-7 cells using a regular imager probe and our fluorogenic probe (Fig. 1F,G). Under these high-density labeling conditions, the limiting factor is the off-rate since multi-emitter artifacts need to be avoided (**Suppl. Fig. 2F**). Regular DNA-PAINT imaging was performed at a frame rate of 4 Hz (expected off-rate ~1 s^−1^) with probe concentration at 0.1 nM, similar to as previously used for high quality DNA-PAINT imaging ^21^. whereas with our fluorogenic imager probe, imaging was performed at 100 Hz (off-rate ~50 s^−1^) and a probe concentration of 20 nM. With both higher on- and off-rates for our fluorogenic probe, approximately 26 times more blinking events per second could be observed than with the regular DNA-PAINT probe (4.3 vs 0.2 s^−1^ µm^−1^ of microtubules; Fig. 1G). Achieving 1,000 blinking events per micron of microtubule required ~4 min of imaging with the fluorogenic probe compared to ~1.4 hr with the regular probe. With the fluorogenic probe, 2D projections of hollow microtubules were easily observable after 3 min (Fig. 1F). Using the regular probe, in contrast, required hours of imaging to obtain similar image quality.

We next tested whether the fluorogenic properties of our new probe would be strong enough to replace the background suppression provided by optical sectioning through TIRF illumination and thereby allow 3D imaging of volumes thicker than a few 100 nm. We imaged microtubules using regular widefield illumination and astigmatic detection for 3D localization. We were able to image with our fluorogenic probe at 10 nM concentration and at a camera frame rate of 100 Hz to acquire a 3D super-resolution image at high quality (FSC resolution = 41.1 nm) in only 10 minutes (Fig. 2 A-C and **Suppl. Fig. 1**). We halved the imager probe concentration compared to the 2D experiments to compensate for the larger size of the blinks caused by the astigmatic detection to avoid an increase in multi-emitter artifacts. The obtained localization precision (**Suppl. Fig. 1C**; peak 1.7 nm for XY, 4.5 nm for Z) was comparable to that reported for regular DNA-PAINT ^21^. The hollow centers of microtubules can be seen in the microtubule cross-section or in the 30-nm thick z-slice as they pass through the plane (Fig. 2B and insert).

Unhindered by the need for optical sectioning, we were also able to perform fast DNA-PAINT under widefield illumination on a 4Pi-SMS microscope that interferometrically combines the fluorescence collected by two opposing objectives for improved axial localization precision ^22, 23^. We imaged mitochondria, immunolabeled for the outer membrane protein OMP25, throughout the cell (Fig. 2D,E; 4 optical sections spanning ~2 µm).

While in this study we demonstrate imaging at 100 fps and generating super-resolution images with excellent localization density and localization precision in minutes, it is worth pointing out that we have not yet reached a fundamental limit with fluorogenic DNA-PAINT. Further developments in probe chemistry and design should yield even better fluorogenicity and faster off-rates. Similarly, on-rates can be further optimized as recently demonstrated for regular DNA-PAINT probes ^11^. Our new probe is as easy to use as the regular probes and is not only compatible with existing DNA-PAINT instruments and modalities, but is now no longer critically dependent on optical sectioning. Importantly, imaging with the fluorogenic probe retains robustness against photobleaching; high signal-to-noise images can be acquired at essentially arbitrarily high sampling densities by extending imaging time (**Suppl. Fig. 1A**). We therefore believe that our new probe concept will become the new gold standard and probe of choice for the majority of DNA-PAINT microscopy.

## Disclosure of Significant Financial Interest

J. B. discloses significant financial interest in Bruker Corp. and Hamamatsu Photonics.

## Acknowledgements

We thank Kevin Hu, Lukas Fuentes and Zach Marin for helpful discussions.

