## Supplementary Material for "Fluorogenic probe for fast 3D whole-cell DNA-PAINT"

### Supplementary Information

### Methods

#### Imager probes and docking strands:

The sequences for the regular imager probe and docking strand were taken from Auer et al., 2017 [15] and share a complementary 9-bases long region. Our fluorogenic probe is 15-bases long, based on an extension of the regular probe. A shortlist of potential imager probe and docking strand sequences was created based on their affinity and lack of secondary structures as predicted by the software BioPython (biopython.org) and NUPACK (nupack.org). The final sequences were selected based on their fluorescence in solution and performance in DNA-PAINT imaging. All oligonucleotides were purchased from Integrated DNA Technologies.

| **Regular DNA-PAINT** | | **5' to 3' direction** |
|---|---|---|
| Imager probe | P00 | Cy3B - AGAAGTAATG |
| Docking strand (antibody) | D00a | Azide - TTTATTACTTCT |
| Docking strand (DNA origami) | D00b | TATTACTTCTT - DNA origami |
| **Fast fluorogenic DNA-PAINT** | | **5' to 3' direction** |
| Imager probe | P10 | Cy3B - AGAAGTAATGTGGAA - BHQ2 |
| Docking full complement for measurements in solution | D10 | TTCCACATTACTTCT |
| Docking strand fast (antibody) | D11a | CCTTCAACATATCCTCTAC - Azide |
| Docking strand fast (DNA origami) | D11b | CCTTCAACATATCCTCTA - DNA origami |

#### DNA origami:

The ring-shaped DNA origami structure is based on a previously described structure [20] with only minor modifications in the positions of handles. The structure was designed using caDNAno (caDNAno.org), with an expected outer diameter of ~62 nm. DNA scaffold strands (8064-nt circular ssDNA) were produced using *E. coli* and M13-derived bacteriophages. All DNA oligonucleotides were purchased from Integrated DNA Technologies.

48 ssDNA handles (docking strands for DNA-PAINT) extend from the 5' end of staple strands on the outer helices (See section above for sequences of the docking strands). Additionally, 12 ssDNA handles extend from the 3' end of staple strands on the bottom of the ring for biotin functionalization.

| | **5' to 3' direction** |
|---|---|
| Biotin handle | DNA origami - CGGTTGTACTGTGACCGATTC |
| Biotin conjugated anti-handle | GAATCGGTCACAGTACAAC - Biotin |

DNA origami rings were assembled from a scaffold strand (80 nM) and a pool of staple strands (480 nM each) in 1× folding buffer (25 mM TrisHCl, 1 mM EDTA, pH 8.0 with 16 mM of $MgCl_2$) using an 18-hr thermal annealing program (85°C–25°C). Folded structures were concentrated by resuspending in half the original volume following PEG precipitation [24]. Correctly assembled rings were purified via rate-zonal centrifugation through 15–45% glycerol gradients (in 1× folding buffer) in an SW 55 rotor (Beckman Coulter) at 48,000 rpm at 4°C for 1.5 hr [25].

DNA origami were attached to channel slides (ibidi USA, µ-Slide VI, 80607) for imaging following a previously published protocol [9] with minor modifications. Slides were plasma-cleaned, coated with biotin-BSA (Millipore Sigma, A8549) and incubated with streptavidin (ThermoFisher, 434302). DNA origami with biotin handles were then attached to the functionalized coverslips. Non-fluorescent beads (Spherotech, TP-08-10) and nanodiamonds (Adámas Nanotechnologies, NDNV140nmHi) were added as fiduciary markers for drift correction.

#### Cell samples:

COS-7 cells were grown in DMEM (Gibco, 21063029) supplemented with 10% fetal bovine serum (FBS; Gibco, 10438026). U-2 OS cells were grown in McCoy's 5A medium (ATCC, 30-2007) supplemented with 10% FBS.

#### Microtubule labeling:

Channel slides (ibidi USA, μ-Slide VI, 80607) were plasma-cleaned and coated with poly-l-lysine (Sigma Aldrich, P4707) before seeding COS-7 cells. Microtubules were labeled following a previously published protocol [23]. COS-7 cells were incubated with 0.2% saponin for 1 min, fixed with 3% paraformaldehyde (Electron Microscopy Sciences, 15710) and 0.1% glutaraldehyde (Electron Microscopy Sciences, 16019) for 15 min, rinsed 3 times with PBS, incubated in blocking buffer (PBS + 0.2% Triton X-100 + 3% bovine serum albumin) for 30 min, and then incubated with mouse anti-alpha tubulin primary antibody (Sigma Aldrich,T5168) at a concentration of 1:200 overnight at 4°C. The cells were then washed 3 times, for 5 min each and incubated with goat anti-mouse IgG secondary antibody (Jackson ImmunoResearch, 115-005-146) at a concentration of 1:200 for 1 hr at room temperature. The secondary antibody was conjugated to oligonucleotide docking strands using Azide / DBCO click chemistry [9]. The cells were washed 3 times for 5 min each and then rinsed 3 times with PBS.

#### Mitochondria labeling:

U-2 OS cells were electroporated with plasmids encoding the mitochondrial marker GFP-OMP25 and then seeded onto ozone-cleaned 25-mm round coverslips. Mitochondria were labeled following a previously described protocol [23]. The cells were fixed with 3% PFA and 0.1% GA for 15 min, permeabilized for 3 min, incubated in blocking buffer for 1 hr, and then labeled with rabbit anti-GFP primary antibody (Invitrogen, A-11122) at 1:500 overnight at 4C. The cells were washed 3 times for 5 min each prior to incubation with goat anti-rabbit IgG secondary antibody (Jackson ImmunoResearch, 111-005-144) at a concentration of 1:200 for 1 hr. The secondary antibody was conjugated to oligonucleotide docking strands using Azide / DBCO click chemistry [9]. The cells were washed 3 times for 5 min each and finally rinsed with PBS 3 times.

#### Fluorescence measurements in solution:

Fluorescence measurements of the probes in solution were performed under a microscope (the same microscope as described for imaging DNA origami structures) under widefield illumination provided by a xenon arc lamp (Sutter Instruments, Lambda LS). Fluorescence intensity was measured ~10 µm deep past the coverslip. Blanks were measured for background correction. Samples were prepared in a high ionic strength PBS-based buffer (PBS, 500 mM NaCl) in channel slides (ibidi USA, µ-Slide VI, 80607). 'Unbound probes' samples contained only the imager probe (0.2 µM), and 'bound probes' samples were prepared with the probe and its complementary sequence in excess (20 µM).

#### Microscope setup:

DNA origami samples and microtubule samples (for 2D) were imaged in TIRF mode on a modified Nikon Ti-E inverted microscope with a 100× 1.45 NA oil immersion objective on a sCMOS camera (Andor, Zyla 4.2). For illumination, a 561-nm laser with a built-in acousto-optic modulator for intensity modulation (Omicron Lux) was used with a dichroic (Semrock, Di02-R488-25x36) and a band pass filter (Semrock, FF01-524/45-25). For imaging with the regular probe, data were recorded at 4 fps at ~0.2 kW/cm$^2$. With the fluorogenic probe, data were recorded at 100 fps at ~2 kW/cm$^2$. A custom image-based focus-lock system was developed based on the a previously described design [26].

Astigmatic 3D imaging of microtubule samples was performed on a custom-built microscope as previously described [27]. Briefly, fluorescent signal was collected by an oil-immersion objective lens (Olympus, 100×, 1.49 NA) and imaged on a sCMOS camera (Hamamatsu, ORCA-Flash 4.0). A cylindrical lens (f = 500 mm) was added to the emission beam path to introduce astigmatism. Data were recorded at 100 fps with a 560-nm laser (MPB Communications, 500 mW) at an intensity of about 13 kW/cm$^2$. 100 nm fluorescent microspheres (ThermoFisher, 580/605, F8801) were imaged to generate reference PSFs. A custom-built focus-lock system based on tracking a reflected infrared laser was used to correct for axial drift.

Mitochondria samples were imaged with a custom-built 4Pi-SMS system as previously described [23]. Briefly, the fluorescent signal was collected coherently by two opposing objectives (100×, 1.35 NA, silicone oil immersion, Olympus) and imaged on a sCMOS camera (ORCA-Flash 4.0v2, Hamamatsu). Data were acquired at 100 fps with a 560 nm laser (2RU-VFL-P-2000-560-B1R, MPB Communications) at an intensity of about 15 $kW/cm^2$.

#### Imaging buffer:

For imaging origami structures, a Tris-based buffer (5 mM Tris, 10 mM $MgCl_2$, 1 mM EDTA, 0.05% Tween 20, 20 mM $Na_2SO_3$ and 1 mM Trolox, pH 7.3-7.5) was prepared. For imaging fixed cell samples, a high ionic strength PBS-based buffer (1× PBS, 500 mM NaCl, 20 mM $Na_2SO_3$ and 1 mM Trolox, pH 7.3-7.5) was used. Trolox (Santa Cruz Biotechnology, sc-200810) aliquots were stored at -20°C at a concentration of 50 mM, and thawed prior to the experiment. Imager probes were stored at -20°C at a concentration of 100 µM in $H_2O$, and serially diluted into one of the imaging buffers as necessary.

#### Data analysis:

Data acquired from the single objective microscope systems were analyzed with PYME (python-microscopy.org) and custom code written in Python (python.org). Localizations in 2D were performed by weighted least square fit with a 2D Gaussian PSF model [28, 29]. Astigmatic 3D localizations were performed by fitting against a PSF experimentally-derived from bead images [30].

The localization routine for 4Pi microscope data has been previously described in detail [23]. Images and movies were rendered using Vutara SRX software (Bruker).

Localizations that appear in consecutive frames (allowing for 1 frame misdetection) at the same position (< 2σ xy-precision) were combined into 'blinks'. This correction takes into account that these localizations do not represent independent samples of the docking strands (See **Suppl. Note 2**). Although this results in fewer localization counts, it avoids artificially inflated blinking rates and overcounting artifacts due to fast camera frame rate and/or slow blinking.

Fine drift corrections were performed using the redundant cross-correlation method [31]. Blinks were rendered as 2/3D Gaussians in images and movies.

Fourier ring correlation (FRC) and Fourier shell correlation (FSC) was computed based on the method described in Nieuwenhuizen, et al., 2013 [32]. A threshold of 0.143 was used. A 5 × 5 × 0.6 µm subregion was used for the FSC calculations presented in **Fig. 2C** and **Suppl. Fig. 1C**.

Analytic study of DNA-PAINT imaging speed and simulations of multi-emitter artifact were performed with code written in Python.

### Supplementary Notes

#### 1. DNA-PAINT background

The greatest weakness of DNA-PAINT is that unbound imager probes contribute large amounts of background fluorescence which can drown out the signal peaks of individual bound probe molecules. The average background level $b$ per area can be estimated as

$$b = \beta \cdot c \cdot \xi \quad \textbf{(Eq. 1)}$$

where $\beta$ is the molecular brightness (i.e. the detected number of photons per second per probe molecule), $c$ is the concentration of imager probes, and $\xi$ is the thickness of the observed volume.

Regular DNA-PAINT relies on (1) low probe concentrations $c$, and (2) optical sectioning, such as total internal reflection fluorescence (TIRF) illumination, to minimize $\xi$ [9]. Regular DNA-PAINT is usually incompatible with widefield illumination due to the large illumination and detection thickness $\xi$ which leads to very high background $b$.

Even when minimizing $\xi$ by optical sectioning, the probe concentration $c$ usually is limited to 5 nM to keep the background fluorescence at an acceptable level [9, 13]. Low concentrations, however, have a detrimental effect on the imaging speed: the blinking rate or binding on-rate $r_{on}$ can be estimated as

$$r_{on} = k_{on} \cdot c \quad \textbf{(Eq. 2)}$$

where $k_{on}$ is the rate constant of an imager probe binding a docking strand.

#### 2. DNA-PAINT imaging speed

The speed of SMLM depends on the time it takes to collect a sufficient number of blinks from every target with the additional caveat that a blink is only useful if it is spatially isolated from other blinks so that it can be correctly identified. Using a binomial distribution model, the probability of detecting $x$ blinks within a diffraction area during an observation period can be predicted from the number of molecular targets (or docking strands; $n$) and the duty cycle ($p$):

$$P(X = x) = \binom{n}{x} \cdot p^x \cdot (1 - p)^{n-x} \qquad \textbf{\textit{(Eq. 3)}}$$

Here, the duty cycle is defined as the ratio of total blinking time ($t_{on}$) for a single emitter and total observation time. For PAINT, it can alternatively be calculated as the binding on-rate ($r_{on}$; proportional to probe concentration) divided by the sum of on- and off ($r_{off}$)-rates:

$$\begin{aligned} p &= \frac{t_{on}}{t_{on} + t_{off}} \\ &= \frac{r_{on}}{r_{on} + r_{off}} \end{aligned} \qquad \textbf{\textit{(Eq. 4)}}$$

With these basic premises, we can investigate the effect of these parameters on imaging duration analytically in order to understand constraints in speed for regular DNA-PAINT and how to overcome them (**Suppl. Fig. 2**).

As a model for SMLM sampling, $P(X = 1)$ estimates the probability of only a single emitter blinking, whereas $P(X > 1)|P(X > 0)$ estimates the fraction of multi-emitter artifacts, i.e. the fraction of blinking events that consists of multiple emitters detected simultaneously within a diffraction-limited volume (**Suppl. Fig. 2C,E**). Assuming each frame represents an observation, then the average imaging duration (number of frames) required to detect a number of blinks equal to the molecular targets is $n/P(X = 1)$. Although we use this here as an estimate of imaging duration, the actual number of frames needed to reliably reconstruct an image is much more complex and beyond the scope of this discussion. The imaging duration is minimized when $P(X = 1)$ is maximized; however, this comes at the cost of a high rate of multi-emitter artifacts. The maximum acceptable multi-emitter artifact rate, for example 10%, presents a threshold and therefore dictates the optimal duty cycle and sets the lower limit for imaging duration.

Since both binding on- and off-rates are tunable in DNA-PAINT, there is an optimal ratio between on- and off-rates to achieve the optimal duty cycle and imaging duration (in frames). However, since image duration in units of time rather than frames is ultimately of interest, it can be added to the model based on the additional assumption that frame rate ($f$) is always matched to binding off-rate:

$$\begin{aligned} T &= \frac{n}{P(X=1)} \cdot \frac{1}{f} \\ &= \left(\frac{r_{on} + r_{off}}{r_{off}}\right)^n \cdot \frac{1}{r_{on}} \end{aligned} \quad \textbf{\textit{(Eq. 5)}}$$

The model predicts that binding on- and off-rates need to be optimized in concert (**Suppl. Fig. 2D,F**; assuming an on-rate constant of $2.3\times10^6$ $M^{-1}$ $s^{-1}$). Increasing the binding on-rate (i.e. by increasing the probe concentration) by itself quickly leads to an unacceptable increase in the rate of multi-emitter artifacts; whereas increasing the binding off-rate alone gains minimal benefits as most of the frames will be blank.

In DNA-PAINT, the maximum acceptable values for both binding on- and off-rates are limited by the signal-to-background ratio (i.e. the ratio between the fluorescence detected from bound vs. unbound probes). To reiterate, the binding on-rate is proportional to probe concentration whereas the blinking off-rate should be proportional to camera frame rate. A fluorogenic probe, by reducing background fluorescence, therefore pushes the acceptable limits of both the binding on- and off-rates allowing the use of higher probe concentrations and higher frame rate.

Our model also recapitulates the effect of different densities of molecular targets $n$. At a fixed blinking off-rate (e.g. the same probe/docking combination), the ideal probe concentration to minimize imaging duration and avoid multi-emitter artifacts decreases with increasing target density (**Suppl. Fig. 2A**). At a fixed probe concentration, higher off-rates are necessary with increasing target density to avoid multi-emitter artifacts (**Suppl. Fig. 2B**). Our imaging experiments, i.e. probe/docking strand affinity and probe concentration, were optimized based on predictions by this model.

### Supplementary Figures

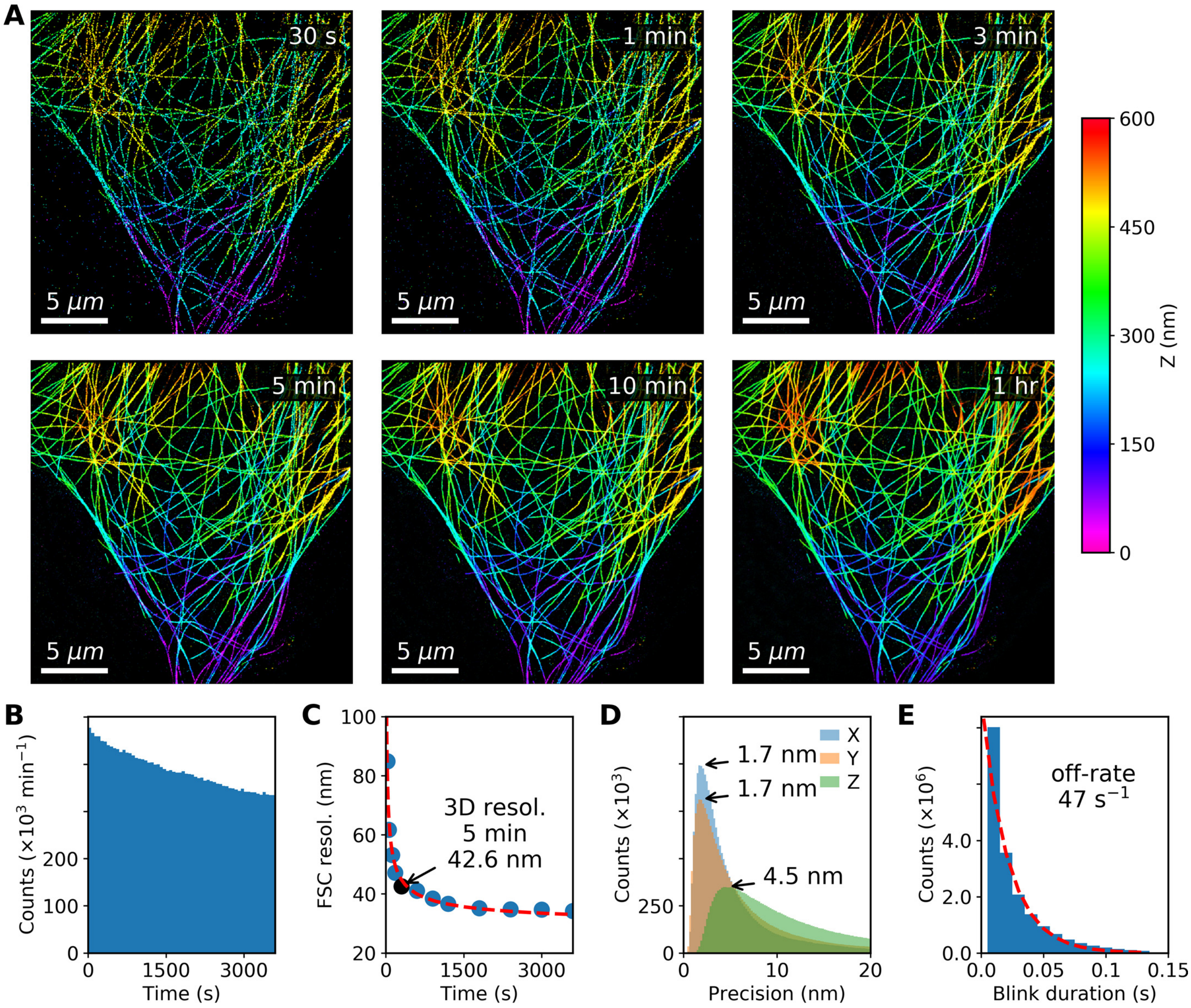

***Supplementary Figure 1***

**Supplementary Figure 1.**

The full dataset from which main **Fig. 2A-C** were generated. **(A)** Fast 3D fluorogenic DNA-PAINT imaging of immunolabeled microtubules in COS-7 cells under widefield illumination at multiple time points. A reasonable low magnification image can be acquired in 30 s. **(B)** Bleaching is negligible, causing only a small reduction (30%) in blinking rate over an hour. **(C)** 3D resolution as quantified by Fourier shell correlation (FSC) improves with longer imaging duration as more blinking events are detected. The resolution reaches 34.3 nm after 1 hr. **(D)** The localization precision peaks at < 5 nm for all three dimensions (X: 1.7 nm, Y: 1.7 nm, Z: 4.5 nm). **(E)** Blink durations fitted with an exponential decay function (blinks that are only 1 frame in duration were ignored for fitting) estimates the mean off-rate at 46.7 $s^{-1}$.

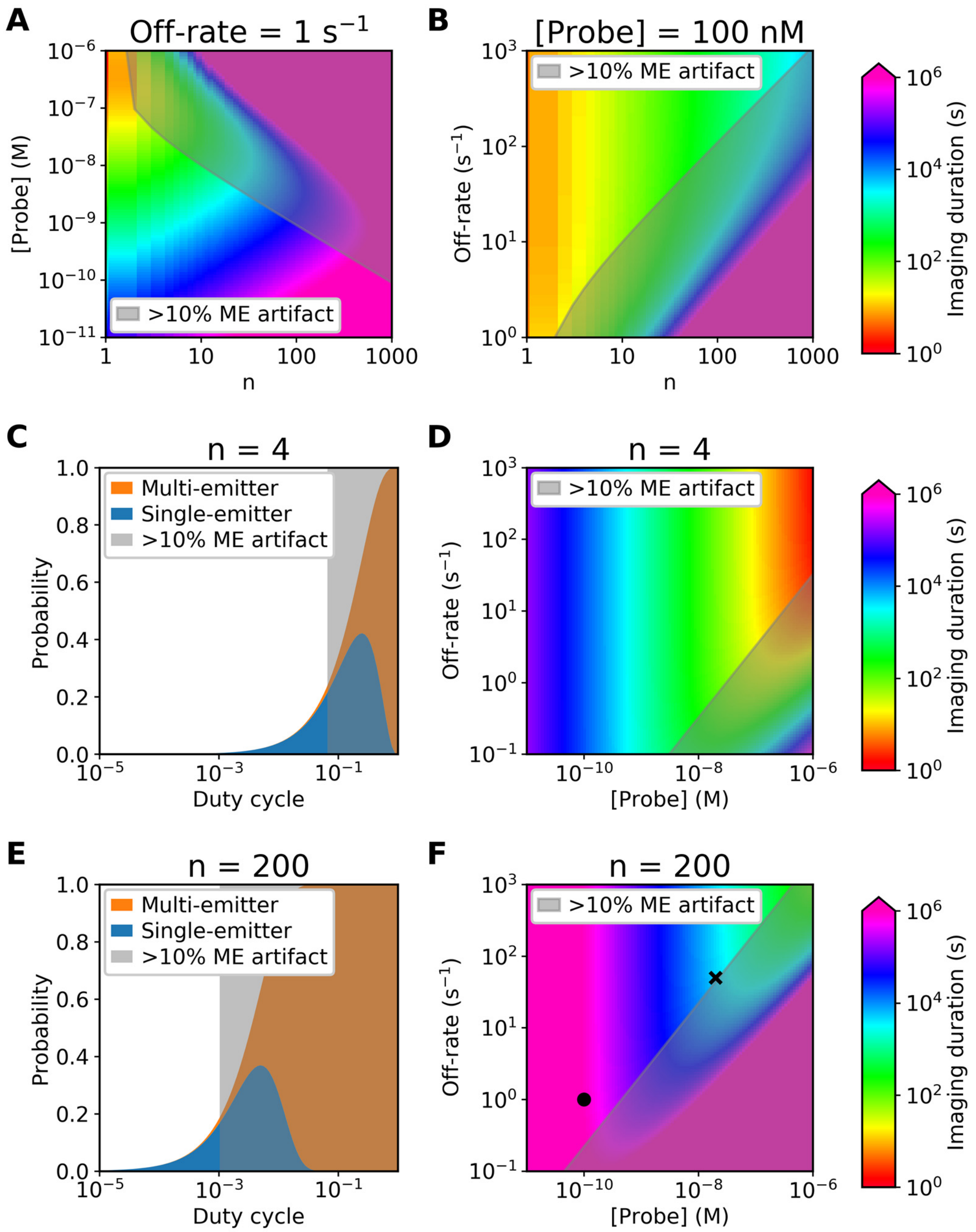

***Supplementary Figure 2***

**Supplementary Figure 2.**

Prediction of DNA-PAINT imaging speed based on a binomial distribution model of blinking. **(A)** For a constant binding off-rate, the optimal imager probe concentration decreases as the number of docking strands in a diffraction-limited area ($n$) increases. **(B)** For a given constant probe concentration (and binding on-rate), the optimal off-rate increases as the number of docking strands in a diffraction-limited area increases. **(C)** A relatively high duty cycle is optimal (6.6%) if there are only $n$=4 docking strands per diffraction-limited area. The upper limit is bound by the need to avoid multi-emitter artifacts. **(D)** Probe concentration and binding off-rate need to be optimized together to minimize imaging duration. For both regular DNA-PAINT (expected off-rate ~1 $s^{-1}$) and our fluorogenic probe (off-rate ~50 $s^{-1}$), the optimal probe concentration of ~30 and ~1500 nM, respectively, is above what we and others have found to be acceptable from a signal-to-background perspective (>5 and >250 nM, respectively). **(E)** A low duty cycle is optimal (0.1%) if there is a high number of docking strands (e.g. $n$=200) per diffraction-limited area (e.g. antibody labeled microtubule). **(F)** At this docking strand density, in contrast to the $n$=4 scenario, the optimal probe concentration of ~0.5 and ~20 nM, respectively, lies well below the limits dictated by background. Imaging speed is instead limited by the 'slow' binding off-rate with this type of samples when performing fast fluorogenic DNA-PAINT. Conditions used for the experiments presented in **Fig. 1F** are indicated (circle: regular DNA-PAINT; cross: fast fluorogenic DNA-PAINT). The imaging speed was increased by increasing both probe concentration and binding off-rate while maintaining multi-emitter artifacts at an acceptable level.

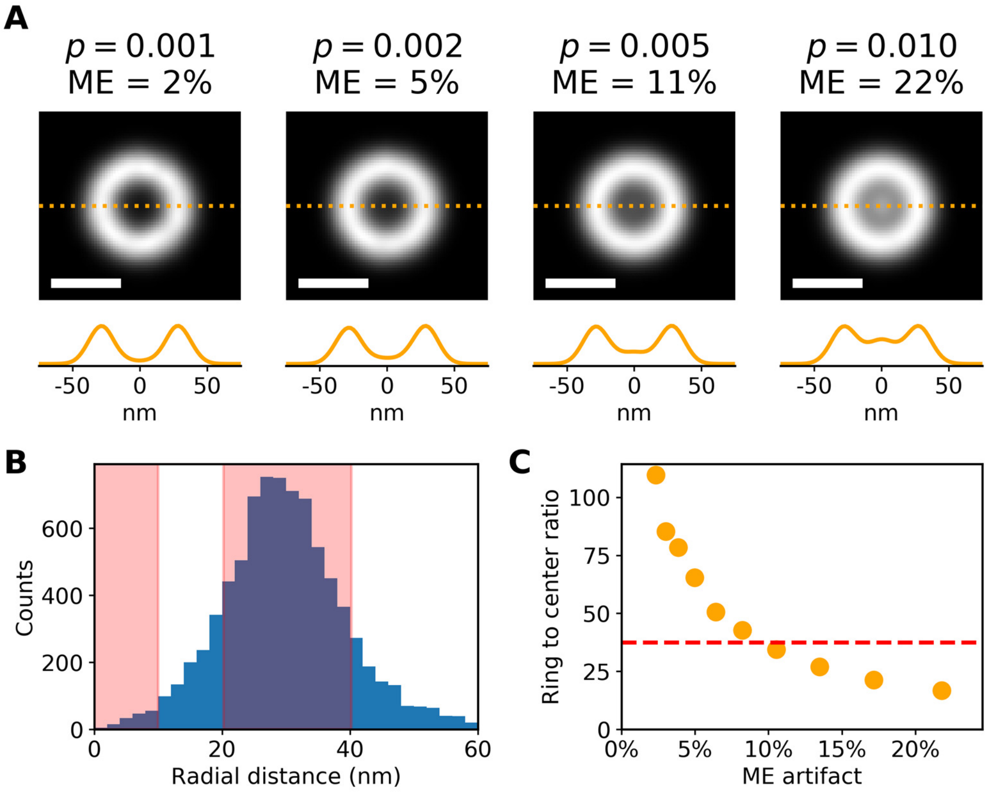

***Supplementary Figure 3***

**Supplementary Figure 3.**

**(A)** Simulation of blinking events on a ring structure (radius = 30.2 nm; docking strands = 48; localization precision = 8.1 nm; observations = 1,000,000) based on a binomial distribution model as described in **Suppl. Note 2**. As duty cycle ($p$) increases, the chance of multi-emitter (ME) artifacts increases which is observed as the erroneous filling in of the ring center. Scale bar is 50 nm. **(B)** This effect was quantified by the ratio of blinking events on the ring (radial distance 20-40 nm; red) versus those in the center (radial distance <10 nm; red). The graph was generated from the origami ring experimental data presented in **Fig. 1D,E**. **(C)** A calibration curve was generated from the simulated data (orange). A ring to center ratio of 37 was calculated from the experimental data (red dash) which estimates the occurrence of multi-emitters at ~9%.
